# Learning an efficient hippocampal place map from entorhinal inputs using non-negative sparse coding

**DOI:** 10.1101/2020.08.12.248534

**Authors:** Yanbo Lian, Anthony N. Burkitt

## Abstract

Cells in the entorhinal cortex (EC) contain rich spatial information and projects strongly to the hippocampus where a cognitive map is supposedly created. These cells range from cells with structured spatial selectivity, such as grid cells in the medial entorhinal cortex (MEC) that are selective to an array of spatial locations that form a hexagonal grid, to weakly spatial cells, such as non-grid cells in the MEC and lateral entorhinal cortex (LEC) that contain spatial information but have no structured spatial selectivity. However, in a small environment, place cells in the hippocampus are generally selective to a single location of the environment, while granule cells in the dentate gyrus of the hippocampus have multiple discrete firing locations but lack spatial periodicity. Given the anatomical connection from the EC to the hippocampus, how the hippocampus retrieves information from upstream EC remains unclear. Here, we propose a unified learning model that can describe the spatial tuning properties of both hippocampal place cells and dentate gyrus granule cells based on non-negative sparse coding from EC input. Sparse coding plays an important role in many cortical areas and is proposed here to have a key role in the hippocampus. Our results show that the hexagonal patterns of MEC grid cells with various orientations, grid spacings and phases are necessary for the model to learn different place cells that efficiently tile the entire spatial environment. However, if there is a lack of diversity in any grid parameters or a lack of hippocampal cells in the network, this will lead to the emergence of hippocampal cells that have multiple firing locations. More surprisingly, the model can also learn hippocampal place cells even when weakly spatial cells, instead of grid cells, are used as the input to the hippocampus. This work suggests that sparse coding may be one of the underlying organising principles for the navigational system of the brain.

**Significance Statement:** The brain can perform extremely complex spatial navigation tasks, but how it does this remains unclear. Here we show that the principle of sparse coding can be used to learn the hippocampal place map in a way that efficiently tiles the entire spatial environment using EC inputs, namely either grid cells or weakly spatial cells. This demonstrates that the hippocampus can retrieve spatial information from the entorhinal cortex using an efficient representation and that sparse coding may be one of the underlying principles of the navigational system of the brain.

## Introduction

Since the Nobel-prize-winning discovery of place cells in the hippocampus (O’Keefe and Dostrovsky, 1971; O’Keefe, 1976) and grid cells in the medial entorhinal cortex (MEC) (Hafting et al., 2005; Rowland et al., 2016), brain regions involved in spatial awareness and navigation have attracted much attention from both experimental and computational neuroscientists.

In the hippocampus, place cells generally have a single specific firing location in a small environment (O’Keefe and Dostrovsky, 1971; Park et al., 2011) and neighboring cells have firing fields at different locations, such that the local cell population in the hippocampus can represent the whole spatial environment (O’Keefe, 1976). In contrast, granule cells in the dentate gyrus of the hippocampal formation have multiple discrete firing locations without spatial periodicity (Jung and McNaughton, 1993; Leutgeb et al., 2007).

However, in the entorhinal cortex (EC), cells that carry spatial information range from cells with structured selectivity to weakly spatial cells that have spatial information but no structured selectivity. For example, MEC grid cells have hexagonal firing fields that cover the entire environment (Hafting et al., 2005). The hexagonal grid is characterised by three grid parameters: spacing, orientation, and phase (See Figure 1A, B and C). Grid cells are organised into discrete modules according to their spacings because the progression in grid spacing along the dorsal-ventral axis is geometric, with ratio around 1.42 (Stensola et al., 2012). Also, grid cells in each module have similar orientation but random phases (Stensola et al., 2012). Nevertheless, there are also other cells such as weakly-tuned cells (Zhang et al., 2013) in the MEC and Diehl et al. (2017) found that, apart from the well-characterised spatial cells in the MEC, nearly all of the remaining 2/3 MEC cells have spatial selectivity. In addition, cells in the lateral entorhinal cortex (LEC) also display weak spatial selectivity (Hargreaves et al., 2005; Yoganarasimha et al., 2011).

**Figure 1:**
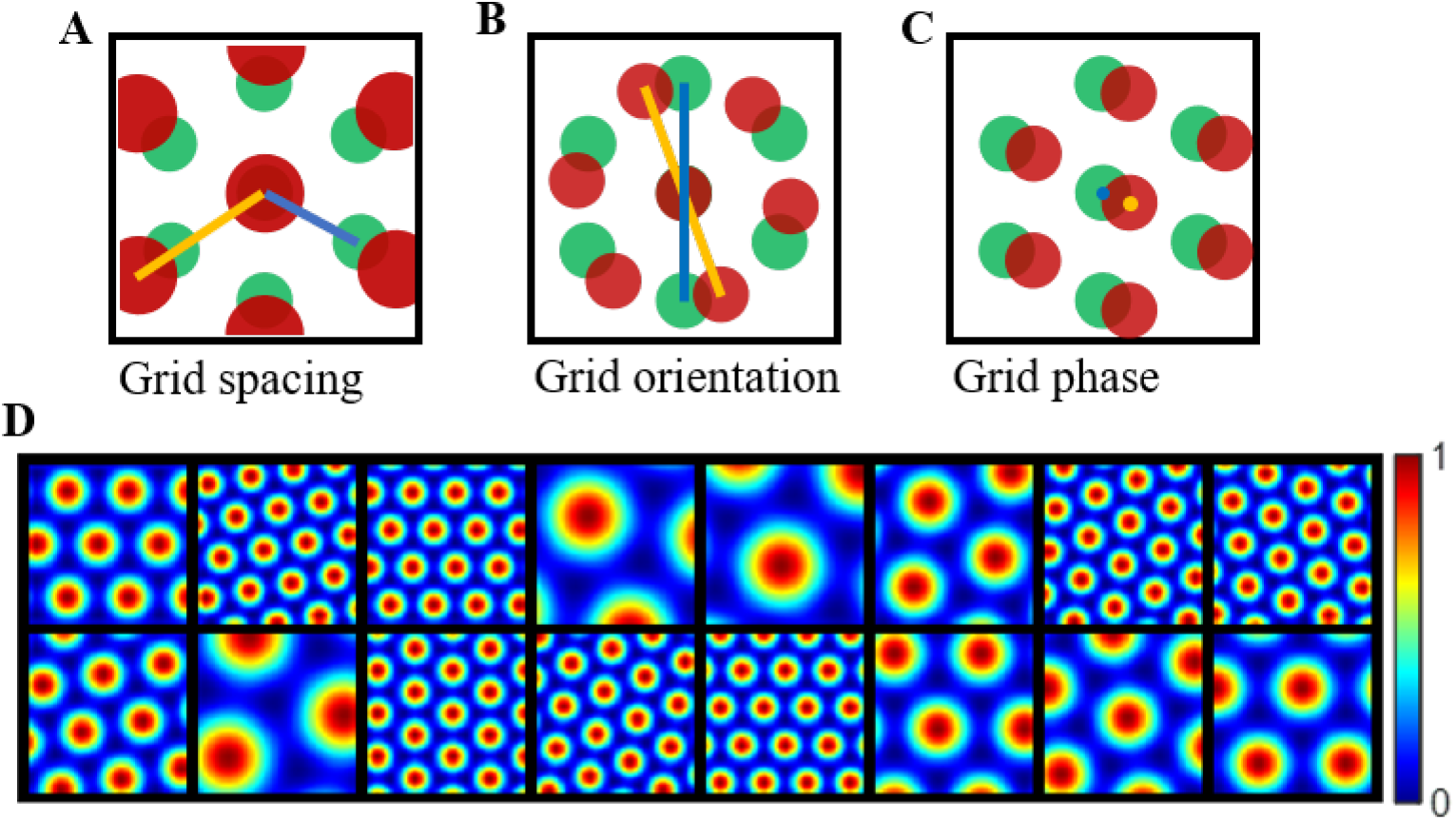
Grid parameters and example grid cells defined by Equation 1. **(A-C)** The schematic plots that illustrate grid spacing, orientation and phase (re-generated from Moser et al. (2014)). **(D)** Grid cells generated by Equation 1 that have different grid spacings, orientations and phases. Each block represents the hexagonal firing field of a grid cell in a 1m × 1m environment. Values in each block are normalised to [0 1] in this plot.

Followed by the Experimental studies that indicates that EC input, including both grid cells and weakly spatial cells, is the principal cortical input to the hippocampus (Steward and Scoville, 1976; Tamamaki and Nojyo, 1993; Leutgeb et al., 2007; Van Strien et al., 2009; Zhang et al., 2013), various models have been proposed to explain the emergence of place cells based on the feedforward connection from grid cells, from mathematical models that have no learning (Solstad et al., 2006; de Almeida et al., 2009) to models with plasticity (Rolls et al., 2006; Franzius et al., 2007b,a; Savelli and Knierim, 2010; Neher et al., 2017).

Among these learning models, Rolls et al. (2006) used a competitive learning procedure. However, only approximately 10% of modelled hippocampal cells had a single firing location. Furthermore, the competition was introduced by manually setting the population activation to a small specified value. Similarly, Franzius et al. (2007a,b) applied independent component analysis (Hyvarinen, 1999) to learn place cells. However, the examples of learned place cells are mostly located at the border (Figure 3C in Franzius et al. (2007a) and Figure 1G in Franzius et al. (2007b)). Additionally, the weights connecting grid and place cells can be positive or negative, and the place cell responses were manually shifted by a constant term to be kept non-negative, which puts into question the biological realisation of the model. Furthermore, previous models do not investigate how well the learned hippocampal place map represents the entire spatial environment and how the weakly spatial cells in the EC can contribute to the formation of the place map.

Sparse coding (Olshausen and Field, 1996) provides a compelling explanation of many experimental findings of brain network structures. One particular variant, non-negative sparse coding (Hoyer, 2003), has recently been shown to account for a wide range of neuronal responses in brain areas (see Beyeler et al. (2019) for a review). However, whether sparse coding can learn the hippocampal place map has not previously been investigated.

Here we applied sparse coding with non-negative constraints, where neuronal responses and connection weights are restricted to be non-negative, to building a learning model of hippocampal cells using EC input. Our results show that, when grid cells are used as the entorhinal input, single-location hippocampal place cells that tile the entire environment can be learned, given the sufficient diversity in grid parameters. However, if there is a lack of diversity in any grid parameters, the learning of the hippocampal place cells is impeded; instead, more hippocampal cells with multiple firing locations are learned. Furthermore, a lower number of hippocampal cells in the network also results in learning hippocampal cells with multiple firing locations. Additionally, the competition generated by sparse coding naturally provides a global inhibition such that learned hippocampal place cells display single firing fields, suggesting that the proposed model can be implemented by biologically based neural mechanisms and circuits. Also, the learned hippocampal place cells tile the entire spatial environment efficiently by a hexagonal lattice, which is consistent with a recent study of conceptual state spaces (Mok and Love, 2019). Moreover, the model can still learn hippocampal place cells even when the grid inputs are replaced by the responses of weakly spatial cells, suggesting that sparse coding can retrieve information from EC input efficiently to form a cognitive map as long as there is sufficient spatial information in the upstream input. This also provides a plausible explanation of many experimental studies that suggest the emergence and maintenance of place cells does not require grid cells (Langston et al., 2010; Wills et al., 2010; Koenig et al., 2011; Brandon et al., 2014; Hales et al., 2014; Schlesiger et al., 2015).

## Materials and Methods

### The environment

The 2D spatial environment used in this study is a 1m × 1m square box. A 32 × 32 grid with 1024 points is used to represent the entire environment. Therefore, a 1024 × 1 vector, denoted by **r**, with only one non-zero element of value 1 can be used to represent the spatial location of a virtual rat.

### Model of entorhinal input

Since the environment is represented by a 32 × 32 grid, a 1024 × 1 vector, denoted by **e**_*i*_, can be used to represent the spatial firing field of the modelled entorhinal cell *i* over the entire environment. For a given position **r** in the environment, the response of modelled entorhinal cell *i* is simply 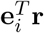. Next, we show how the entorhinal input, grid cells or weakly spatial cells, are modelled to provide information for modelled hippocampal cells.

#### Grid cells described by a mathematical model

The hexagonal firing fields of grid cells can be represented by the sum of three sinusoidal gratings (Solstad et al., 2006; Kropff and Treves, 2008; de Almeida et al., 2009), as described by

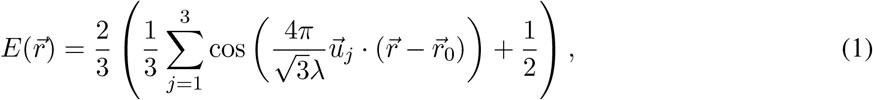

where 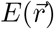 is the grid cell response at the spatial location 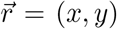 is the grid spacing, *θ* is the grid orientation, 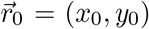 represents the phase offset, and 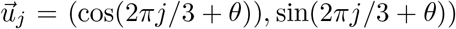 is the unit vector with direction 2*πj/*3 + *θ. E*(*·*), described in Equation 1, is normalised to have a maximal value of 1 and minimum of 0. Because of the periodicity of the hexagonal pattern, the grid orientation, *θ*, lies in the interval of [0, *π/*3), and the phases in both *x* and *y* axes are smaller than the grid spacing, i.e., 0 *≤ x*_0_, *y*_0_ *< λ*.

Since grid cells have different spacings, orientations and phases, Equation 1 is used to generate diverse grid cells. The value of the grid spacing, *λ*, ranges in value from 28 cm (Hafting et al., 2005; Solstad et al., 2006) and increases by a geometric ratio 1.42 that is consistent with experimental results (Stensola et al., 2012) and the optimal grid scale derived by a mathematical study (Wei et al., 2015). For example, if there are *N*_*λ*_ different grid spacings, the spacings will be 28 cm, 28 × 1.42 = 39.76 cm, *…*, and 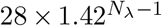. For each grid spacing, different values of grid orientation, *θ*, are uniformly taken from the interval [0, 60°). For example, if there are 3 different grid orientations, the values will be 0, 20° and 40°. The number of different orientations for each grid spacing is denoted as *N*_*θ*_. Furthermore, it is assumed here that there are *N*_*x*_ and *N*_*y*_ phases along x-axis and y-axis for each specific grid spacing and orientation. Similar to grid orientation, the value of the phase is taken uniformly from [0, *λ*). For example, if there are 2 different phases along the x-axis, they will have the values *x*_0_ = 0 and *λ/*2. The resulting total number of modelled entorhinal cells (grid cells), denoted as *N*_e_, will be the product of numbers of spacings, orientations and phases:

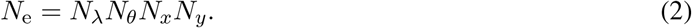

Some examples of grid cells described by Equation 1 are shown in Figure 1D. These grid cells have diverse grid spacings, orientations and phases.

The number of different grid parameters (*N*_*λ*_, *N*_*θ*_, *N*_*x*_ and *N*_*y*_) of grid cells defined above are assigned to different values to investigate the effect of the diversity of grid cells on the formation of place cells.

Though the firing fields of grid cells can be represented by the sum of three sinusoidal gratings (Equation 1) given grid spacings, orientations, and phases and are used in many studies (Solstad et al., 2006; Kropff and Treves, 2008; de Almeida et al., 2009), this mathematical model cannot capture the variability in individual grid fields of the grid cells as reported by Ismakov et al. (2017). Therefore, in this paper, we also adopt a more realistic model of grid cells that characterises grid fields individually and embraces the variability between individual grid fields. This model of grid cells is used to investigate the robustness of our results obtained by using the mathematical model of grid cells (Equation 1).

#### Grid cells described by a more realistic model

MEC Grid cells have different spacings, orientations and phases, and are separated into discrete modules where grid cells are anatomically adjacent and have similar grid spacings, which is supported by experimental evidence (Hafting et al., 2005; Stensola et al., 2012). Moreover, grid cells in the same spacing module tend to have similar orientations and random phases (Stensola et al., 2012). Because of these properties of grid cells, we model grid cells using a similar approach in a previous study (Neher et al., 2017) as described below.

Values of the grid spacing in each module are randomly sampled from normal distributions: there are four discrete modules for grid spacings (*λ*) with mean of 38.8 cm, 48.4 cm, 65 cm and 98.4 cm, and a same standard deviation of 8 cm. For grid orientation (*θ*), since grid cells in the same spacing module tend to have similar orientations, grid cells in the four discrete modules also have mean orientations 15°, 30°, 45° and 0°, and a common standard deviation of 3°. Because grid phases, (*x*_0_, *y*_0_), are random in each module, grid phase is randomly sampled from a uniform distribution between 0 and *λ*. Stensola et al. (2012) showed that 87% of grid cells belong to the two modules with small spacings based on their recordings. Therefore, we have 43.5%, 43.5%, 6.5% and 6.5% of grid cells in the modules with mean spacings 38.8 cm, 48.4 cm, 65 cm and 98.4 cm, respectively (unless otherwise noted).

The firing field of each grid cell is modeled as the sum of multiple grid fields whose centers are located at the vertices of the hexagonal grid. The grid field at vertex (*x*_*v*_, *y*_*v*_) is described by a function with the following form (Neher et al., 2017)

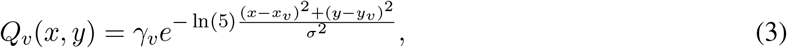

where *γ*_*v*_ is the amplitude, *σ* determines the radius of the grid field, and the response will be *γ*_*v*_/5 at a distance *σ* away from the center. *σ* is determined by the grid spacing, *λ*, with *σ* = 0.32*λ* (Neher et al., 2017). To incorporate the variability of individual grid fields, the amplitude of the grid field at every vertex of the hexagonal pattern, *γ*_*v*_, is chosen from a normal distribution with mean 1 and standard deviation 0.1 (Neher et al., 2017). The locations of all vertices of the hexagonal grid are determined by grid spacing, *λ*, grid orientation, *θ*, and grid phase, (*x*_0_, *y*_0_).

After the mathematical model of grid cells are used as the input, the more realistic model of grid cells are then used to verify the results and investigate the robustness of the model.

#### Weakly spatial cells

Apart from grid cells in the MEC, other cells such as weakly-tuned cells (Zhang et al., 2013) and non-grid cells (Diehl et al., 2017) in the MEC, and LEC cells that contains spatial information (Hargreaves et al., 2005; Yoganarasimha et al., 2011), can also contribute to the formation of hippocampal place cells. In this paper, we model these cells as *weakly spatial cells*. The firing field of weakly spatial cells is generated in the simulation by first assigning a random activation, sampled from a uniform distribution between 0 and 1, to each location, then smoothing the map with a Gaussian kernel with standard deviation 6 cm, and normalising the map such that the values are between 0 and 1 (Neher et al., 2017).

Weakly spatial cells are used as another type of entorhinal input to investigate how they contribute to the formation of the hippocampal place map.

### Sparse coding with non-negative constraints

**Sparse coding** was originally proposed by Olshausen and Field (1996) to demonstrate that simple cells in the primary visual cortex represent their sensory input using an efficient neuronal representation, namely that their firing rates in response to natural images tend to be sparse (rarely attain large values) and statistically independent. In addition, sparse coding finds a reconstruction of the sensory input through a linear representation of features with minimal error, which can be understood as minimising the following cost function

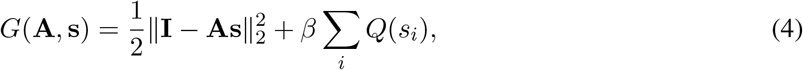

where the matrix **I** is the input, columns of **A** are basis vectors (universal features) from which any input can be constructed from a weighted sum, the vector **s** represents the neural responses and each element, *s*_*i*_, is the coefficient for the corresponding basis vector, the function *Q*(·) is a function that penalises high activity of model units, and *β* is a sparsity constant that scales the penalty function (Olshausen and Field, 1996, 1997). Implemented in a neural network, **A** represents the connection between layers and **s** represents the neuronal responses. The term **As** in Equation 4 represents the model reconstruction of the input, so this cost function represents the sum of squared reconstruction error and response penalty. Therefore, the model finds a sparse representation for the input by solving this minimization problem. By taking the partial derivatives of Equation 4 in terms of the elements of **A** and **s**, and then applying gradient descent, the dynamic equation and learning rule are given by

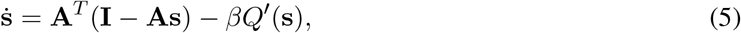

and

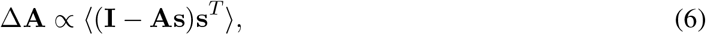

respectively. *Q′*(·) is the derivative of *Q*(·), the dot notation represents differentiation with regard to time, and ⟨ · ⟩ is the average operation.

One common choice of *Q*(·) is the *l*_1_ norm, i.e. the absolute value function. In this case, Rozell et al. (2008) shows that the dynamics in Equation 5 can be implemented via thresholding and local competition in neural circuits, as described by

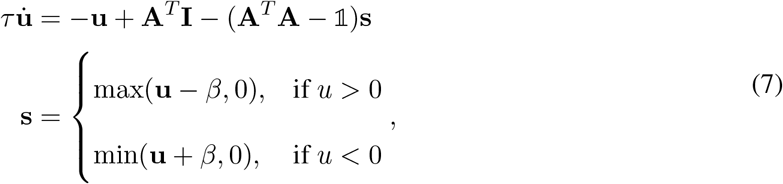

where 1 is the identity matrix, *τ* is the time constant, **u** can be interpreted as the membrane potential, and *β* is the positive sparsity constant in Equations 4 and 5 and becomes the threshold of the thresholding function in Equation 7. In this paper, we will use Equation 7 (Rozell et al., 2008) to implement sparse coding and use the learning rule in Equation 6 to update the entorhinal-hippocampal connection **A**.

**Non-negative sparse coding** is simply sparse coding with non-negative constraints, i.e., the connection weights **A** and model responses **s** are restricted to non-negative values in the cost function Equation 4. Note that, when *β* in Equation 4 is set to zero, the cost function of non-negative sparse coding reduces to the cost function of non-negative matrix factorization (Lee and Seung, 1999).

### Structure of the model

In this study, a two-layer network is proposed to model the activities of entorhinal cells (first layer) and hippocampal cells (second layer), respectively. Given a spatial location in the environment, modelled en-torhinal cells respond according to their firing fields. Modelled entorhinal cell responses then feed into modelled hippocampal cells and the entorhinal-hippocampal network implements a sparse coding model with non-negative constraints. The model structure is shown in Figure 2.

**Figure 2:**
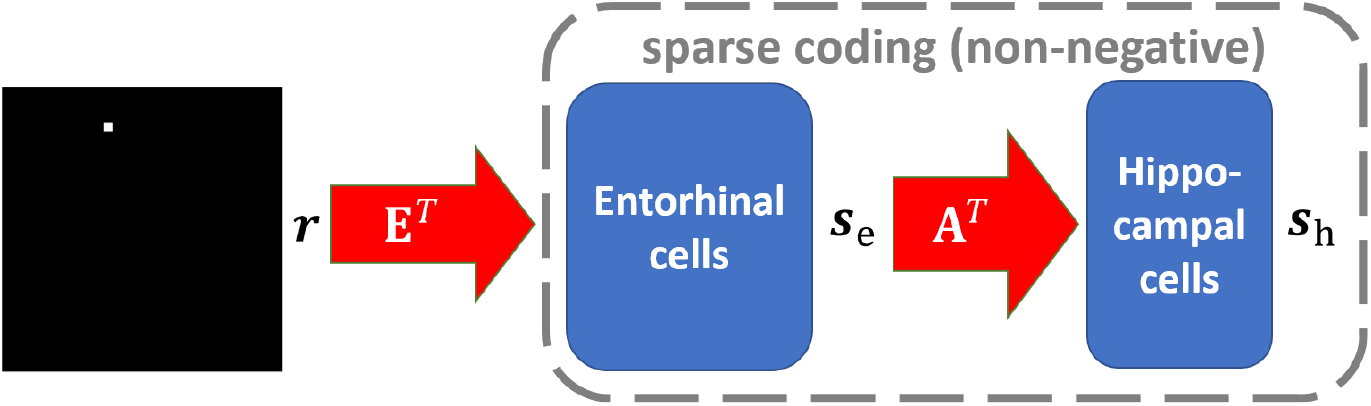
Graphical representation of the model. Red arrows represent non-negative connections. Notation is defined in the main text. **E** represents the firing fields of modelled entorhinal cells and **A** represents the connection between the entorhinal cortex and hippocampus.

Denote **E** as a 1024 × *N*_e_ matrix that represents the firing fields for *N*_e_ modelled entorhinal cells in the network; i.e. each column of **E, e**_*i*_ (*i* = 1, 2, …, *N*_e_), is a 1024 × 1 vector that represents the firing field of modelled entorhinal cell *i*. For a spatial location **r** in the environment, modelled entorhinal cell responses (firing rates), **s**_e_, are given by **s**_e_ = **E**^*T*^ **r**. Modelled hippocampal cell responses (firing rates), **s**_h_, are computed by a sparse coding model for the entorhinal-hippocampal network with non-negative connection **A**. Assume there are *N*_h_ modelled hippocampal cells in the network. Then **A** is a *N*_e_ × *N*_h_ matrix and **s**_h_ is a *N*_h_ ×1 vector. Denote **u**_h_ as a *N*_h_ ×1 vector that represents membrane potentials of modelled hippocampal cells. Based on Equation 7, the dynamics of the model is given by

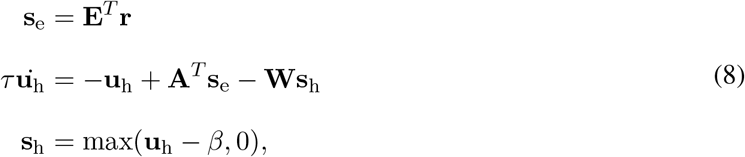

where *τ* is the time constant for modelled hippocampal cells, *β* is the threshold of the rectifying function of firing rates, and **W** can be interpreted as the matrix of recurrent connections between modelled hippocampal cells determined by **W** = **A**^*T*^ **A** − 𝟙, where 𝟙 is a *N*_h_ × *N*_h_ identity matrix. The dynamics of modelled hippocampal cells described in Equation 8 is derived from the local competitive algorithm (LCA) proposed by Rozell et al. (2008) that solves sparse coding efficiently. However, modelled hippocampal cell responses, **s**_h_, and connection matrix, **A**, are taken to be non-negative in this study.

The code to run the model is available at https://github.com/lianyunke/Learning-Hippocampal-Cells-from-EC-Cells-Using-Nonnegative-Sparse-Coding.

### Learning rule

The learning rule for updating the connection strength matrix **A** is similar to that in previous studies of sparse coding (Olshausen and Field, 1997; Zhu and Rozell, 2013), as given by

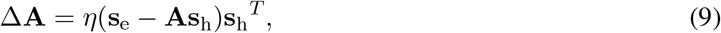

where *η* is the learning rate. Elements of **A** are kept non-negative during training, i.e., the element will be set to 0 if it becomes negative after applying the learning rule described in Equation 9. Then each column of **A** is normalised to unit length, similar to previous studies (Olshausen and Field, 1997; Rolls et al., 2006; Zhu and Rozell, 2013; Lian et al., 2019).

The model dynamics and learning rule described in Equations 8 and 9 can be implemented in a biologically realistic network (Lian et al., 2019). Here we simply use the equations described above to demonstrate that the principle of non-negative sparse coding can learn hippocampal cells with both single and multiple firing locations.

### Training

For modelled entorhinal cells, when using grid cells described by a mathematical model (Equation 1 in Materials and Methods) as the entorhinal input, the smallest grid spacing and grid ratio are taken to be 28 cm and 1.42, respectively. Since the environment used in this study is 1m × 1m, the maximal grid spacing is taken to be smaller than 1m, which leads to 1 *≤ N*_*λ*_ *≤* 4. Therefore, all possible grid spacings are 28 cm, 39.76 cm, 56.46 cm and 80.17 cm. For grid orientation, we have 1 *≤ N*_*θ*_ *≤* 7. For grid phase, we have the same number of phases in each direction and the maximal number is 5, i.e. 1 *≤ N*_*x*_ = *N*_*y*_ *≤* 5. After investigating the effect of grid diversity on modelled hippocampal cells for the learning model, more realistic grid cells (defined in Materials and Methods) are then used to investigate the robustness of the model. Furthermore, weakly spatial cells (defined in Materials and Methods) are also used as the entorhinal input in order to investigate what weakly spatial cells in the EC contribute to place cells’ firing. For both cases of realistic grid cells and weakly spatial cells, the number of modelled entorhinal cells is taken to be 600 (*N*_e_ = 600).

There are 100 modelled hippocampal cells at the second layer in our simulations (unless otherwise noted), i.e., *N*_h_ = 100. The dynamical system described by Equation 8 is implemented by the first-order Euler method, where the membrane time constant is *τ* = 10ms, consistent with the physiological value (Dayan and Abbott, 2001), the threshold is *β* = 0.3, and there are 200 integration time steps with a time step of 0.8ms which we found to provide numerically stable solutions. We use 20, 000 epochs in our training. In each epoch, a random location, **r**, is presented to the grid cells and the model responses are computed using Equation 8 and the matrix of connection strengths, **A**, is updated by Equation 9. The learning rate, *η*, is\ chosen to be 0.03. The parameters above were chosen to ensure a stable solution in a reasonable time scale, but the results were found to be robust to moderate changes of these parameters.

### Recovering the firing fields of modelled hippocampal cells

Because the response of modelled hippocampal cells is computed by the dynamic equations in Equation 8, **A**^*T*^ **s**_e_ = (**EA**)^*T*^ **r** cannot simply represent the response corresponding to the spatial location **r** due to the recurrent connection **W**; i.e., **EA** cannot represent the firing field of modelled hippocampal cells. Therefore, after training, we use the method of reverse correlation to recover the firing fields, denoted as **F**, of modelled hippocampal cells. We present *K* uniformly sampled random locations, **r**_1_, *…*, **r**_*K*_, to the model, compute according to Equation 8 the neural responses of a modelled hippocampal cell, *s*_1_, *…, s*_*K*_, and then compute the firing field, **F**, of this modelled hippocampal cell by

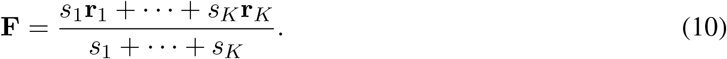

*K* = 10^5^ is used in this paper.

#### Fitting firing fields to functions

In order to obtain the center and size of the firing field of the modelled hippocampal cell, **F** is fitted by a function *Q*(*x, y*) of the form

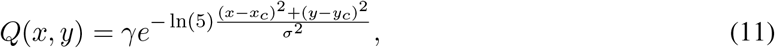

where *γ* is the amplitude, *σ* is the breadth of the firing field, and (*x*_*c*_, *y*_*c*_) represents the center of the function. The built-in MATLAB (version R2020a) function, *lsqcurvefit*, is used to fit these parameters. The fitting error is defined as the square of the ratio between the fitting residual and firing field. After fitting, the fitting error and *σ* are used to determine whether a modelled hippocampal cell meets the criteria of place cell.

#### Selecting place cells

Some firing fields of modelled hippocampal cells have multiple firing locations and noise in the background, while others have a single location of firing. After fitting firing fields into the function described in Equation 11, a modelled hippocampal cell is categorised as a *place cell* if the following two criteria are satisfied for the firing field: (1) the fitting error is smaller than 15% (2) the breadth, *σ*, is larger than 5 cm. These two rules exclude any modelled hippocampal cells with no obvious firing field or with multiple-location firing field. The firing field of a place cell is called *place field*.

### Measuring the uniformity of place cell representation

For place cells that meets the criteria defined above, the field center (*x*_*c*_, *y*_*c*_) fitted by Equation 11 indicates the spatial location that the place cell responds to. We measured how well all place cells represent the entire environment using two measures.

The first measure is *distance to place field, d*_PF_, which indicates the Euclidean distance between each spatial location (*p*_*x*_, *p*_*y*_) in the environment and the nearest place field, described as

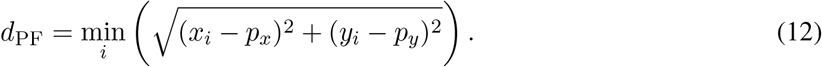

If the distance to a place field is large for a location, it means that there are no place fields near this location. Therefore, the distribution of this measure can tell us how well place fields of all place cells tile the entire spatial environment. When all spatial locations have small values of *d*_PF_, the entire environment is tiled by the place cells.

The second measure is *nearest distance, d*_ND_. We define *d*_ND_ of place cell *j* as the maximal Euclidean distance of 2 nearest centers, described as

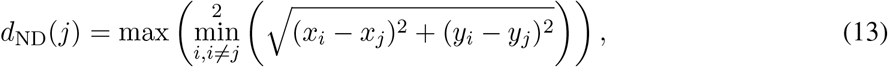

where (*x*_*j*_, *y*_*j*_) is the center of the place field, min^2^ returns a set of 2 smallest values. The distribution of *d*_ND_ for all centers shows the uniformity of place cells in the environment. However, this measure alone provides little information about the coverage of all place cells because place cells with small nearest distance might only lie in a small sub-region of the entire environment, which would give a small value of this measure but would not represent a good tiling of the entire environment.

The distance to place field, *d*_PF_, together with nearest distance, *d*_ND_, provide quantitative measures of how well the place cells code for the spatial environment. Small values of both measures indicate that place cells can tile the entire environment fairly evenly. For example, if 100 place cells are organised on a 10 × 10 grid that evenly tile the 1m × 1m environment, the nearest distance will be 100/(10 − 1) *≈* 11.11 cm for each place cell and the distance to place field for every location is smaller than 11.11/2 *≈* 5.56 cm.

## Results

### When diverse grid cells are used as the entorhinal input, the model can learn place cells that tile the entire environment efficiently

The results shown in this section use grid cells described by the mathematical model as the entorhinal input. Our simulation shows that the non-negative sparse coding model proposed here can learn single-location place cells given diverse grid cells as the input. Grid cells with 4 different spacings (*N*_*λ*_ = 4), 6 different orientations (*N*_*θ*_ = 6) and 5 different phases in both *x* and *y* axes (*N*_*x*_ = *N*_*y*_ = 5) are used here, i.e., there are 600 grid cells in total used as the entorhinal input (*N*_e_ = *N*_*λ*_*N*_*θ*_*N*_*x*_*N*_*y*_ = 600).

After learning, all 100 modelled hippocampal cells meet the criteria of a place cell (as defined in Materials and Methods). The firing fields, **F** (defined by Equation 10), for 100 modelled hippocampal cells are shown in Figure 3A. The firing fields of modelled hippocampal cells are ordered by their spatial locations. All place cells have a single-location firing field. Furthermore, different cells have spatially different firing fields.

**Figure 3:**
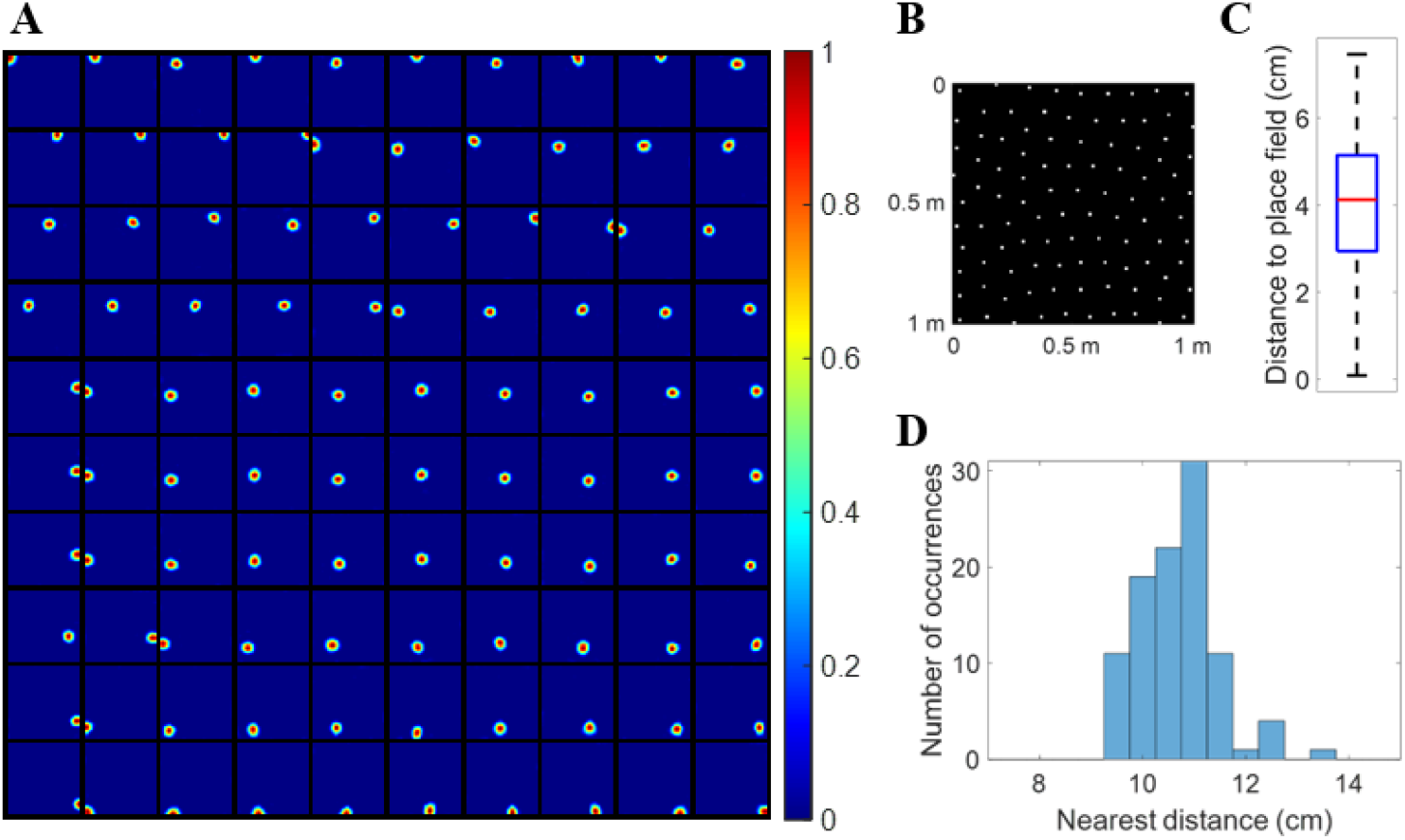
Firing fields and their centers of place cells learned by the model with *N*_*λ*_ = 4, *N*_*θ*_ = 6 and *N*_*x*_ = *N*_*y*_ = 5. **(A)** Firing fields, **F** (defined by Equation 10), for 100 place cells. All 100 modelled hippocampal cells learn a place field at different spatial locations and meet the criteria of a place cell. Each block represents the firing field of a cell in a 1m × 1m environment. Values in each block are normalised to [0 1]. The radii of all place cells have mean 8.92 cm and standard deviation 0.49 cm. **(B)** Centers of the place cells plotted together in the environment, showing that they evenly tile the whole spatial environment with a hexagonal pattern. **(C)** Box plot of distance to place field for all spatial locations in the spatial environment. The black lines at the bottom and top indicate the minimum and maximum, and the bottom edge of the blue box, red line inside the blue box and top edge of the blue box represent 25%, 50% (median) and 75% percentile of the data. In this box plot there are no outliers. **(D)** Histogram of nearest distance for all place cells.

#### The learned place cells tile the environment efficiently

The centers of all the place cells are displayed together in the 1m × 1m spatial environment represented by a 32 × 32 pixel-like image, Figure 3B, which shows that the centers of the 100 place cells tile the entire environment without any overlap. In addition, the box plot in Figure 3C shows that any location within the space is within a distance of no more than 8.2 cm from the nearest place fields. The histogram of nearest distance of all 100 place cells is displayed in Figure 3D, which shows that the distribution is centered around a mean value of 10.70 cm and standard deviation 0.75 cm. Given that the learned place cells have mean radius 8.92 cm with standard deviation 0.49 cm, Figure 3B, C and D illustrate that the learned place cells tile the whole environment rather evenly, i.e, the model learned by non-negative sparse coding can give an accurate neural representation of the spatial location in the environment.

Moreover, centers of all learned place cells (Figure 3B) are positioned into a hexagonal pattern, which can be explained by the principle of the model. Because our model is based on sparse coding that finds an efficient representation of the input, the fact that all learned place cells altogether form a hexagonal pattern is efficient because the triangular lattice of the hexagonal pattern is known to be the optimal solution to the circle packing problem (Thue, 1892). The recurrent connection **W**, described in Equation 8 might provide the inhibition needed for generating an efficient population code of the space. This result is also consistent with a cluster learning method of concept learning in both spatial and conceptual domains (Mok and Love, 2019), suggesting that sparse coding might be an underlying principle of processing conceptual information as well.

#### The competition introduced by sparse coding provides the inhibition for place cells

The connectivity profile between 600 grid cells and 100 place cells is plotted in Figure 4A, which shows that each place cell selects a group of particular grid cells with different weights. As a result, the overall feedforward connection from the spatial environment to the place cells, namely the matrix product **EA**, has the spatial structure plotted in Figure 4B, which shows that each place cell is selective to one spatial location similar to the firing fields (Figure 3A). However, **EA** has strong average offsets, which can be seen from the grey background in Figure 4B. The model of place cells proposed by Solstad et al. (2006) has an inhibition term to balance the excitation so that the place fields are responsive to a single location. As for the model of place cells proposed by Franzius et al. (2007b), an offset constant is added and signs of model units are adjusted in order to achieve single location place fields. Nevertheless, comparing Figure 3A and Figure 4B, we can conclude that the network implemented by sparse coding naturally introduces the competition to inhibit place cells such that they have firing fields similar to those found in experiments. As stated earlier in Materials and Methods, the sparse coding model used in this paper can be implemented by a biologically realistic network (Lian et al., 2019), suggesting that principle used here can be a potential mechanism used in the navigational system of the brain. The results presented in this study are not sensitive to different parameter values as long as there are diversities in spacing, orientation and phase. Even 81 grid cells (*N*_*λ*_ = 3, *N*_*θ*_ = 3 and *N*_*x*_ = *N*_*y*_ = 3) are sufficient for the model to learn place cells that tile the whole environment.

**Figure 4:**
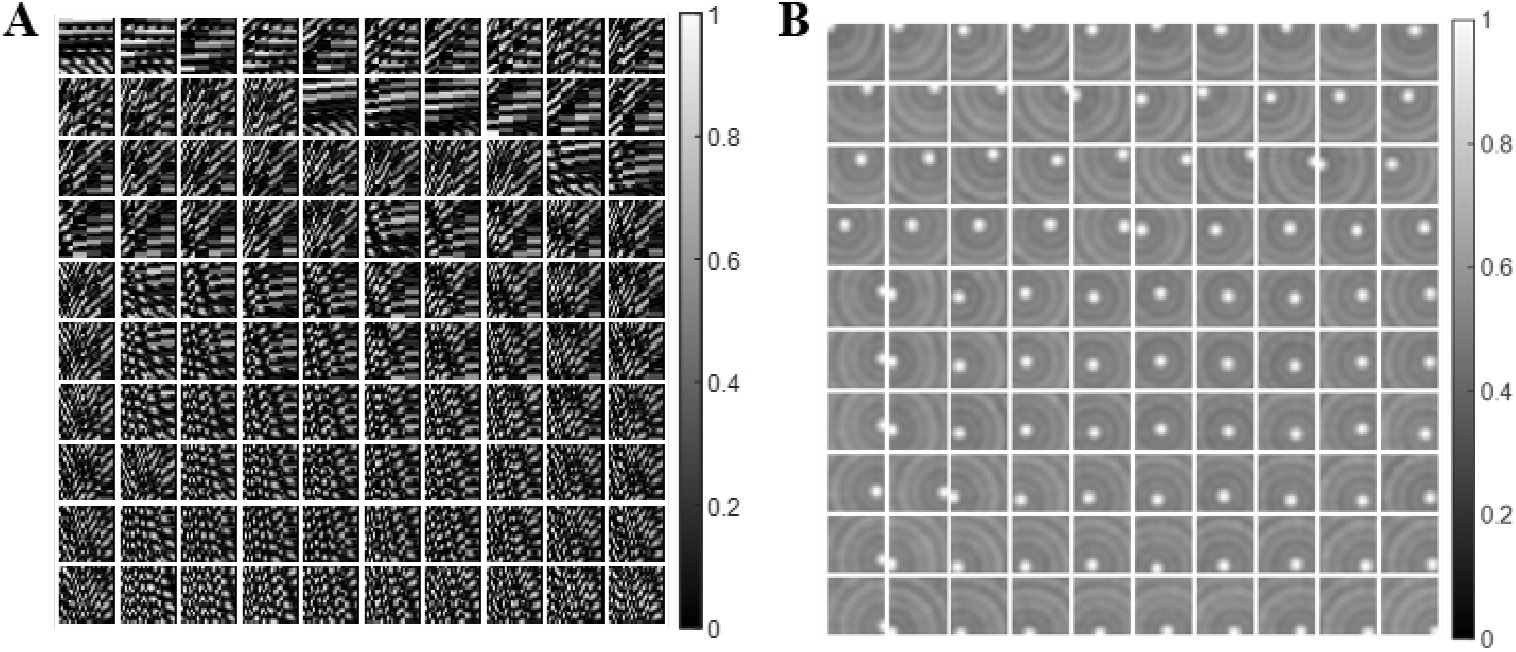
Feedforward connections of the model. **(A)** Connection strengths between grid and place cells, **A**. Each block represents the connections between 600 grid cells and a place cell on a 24 × 25 matrix. **(B)** Feedforward connection strengths from the spatial environment to place cells, **EA**. Each block shows the selective spatial structure of a place cell. For both (A) and (B), values in each block are normalised to [0, 1] for plotting.

In addition, the principle of sparse coding forces the model to learn an efficient representation of the entorhinal input. The average percentage of active modelled hippocampal cells in response to a spatial location is 5.59%. The sparse population activity is consistent with the experimental study that shows sparse ensemble activities in the macaque hippocampus (Skaggs et al., 2007).

### The model can learn cells with multiple firing locations

In this section, we still use grid cells described by the mathematical model as the entorhinal input and show that the lack of diversity in any grid parameters or the lack of modelled hippocampal cells in the network will prevent the model from learning place cells and cells with multiple firing locations start to emerge, i.e., the same model can learn cells similar to dentate gyrus cells that have multiple firing locations.

#### When grid cells are less diverse

A lack of diversity in grid spacing results in the emergence of multiple firing locations of the modelled hippocampal cells, as illustrated in Figure 5A compared with Figure 3A. Similarly, compared with Figure 3B and C, the lack of diversity in grid orientation or grid phase will also cause the model to learn more cells with multiple firing locations (Figure 5B and C). These modelled hippocampal cells are similar to dentate gyrus cells that are found to have multiple firing locations in experimental studies (Jung and McNaughton, 1993; Leutgeb et al., 2007).

**Figure 5:**
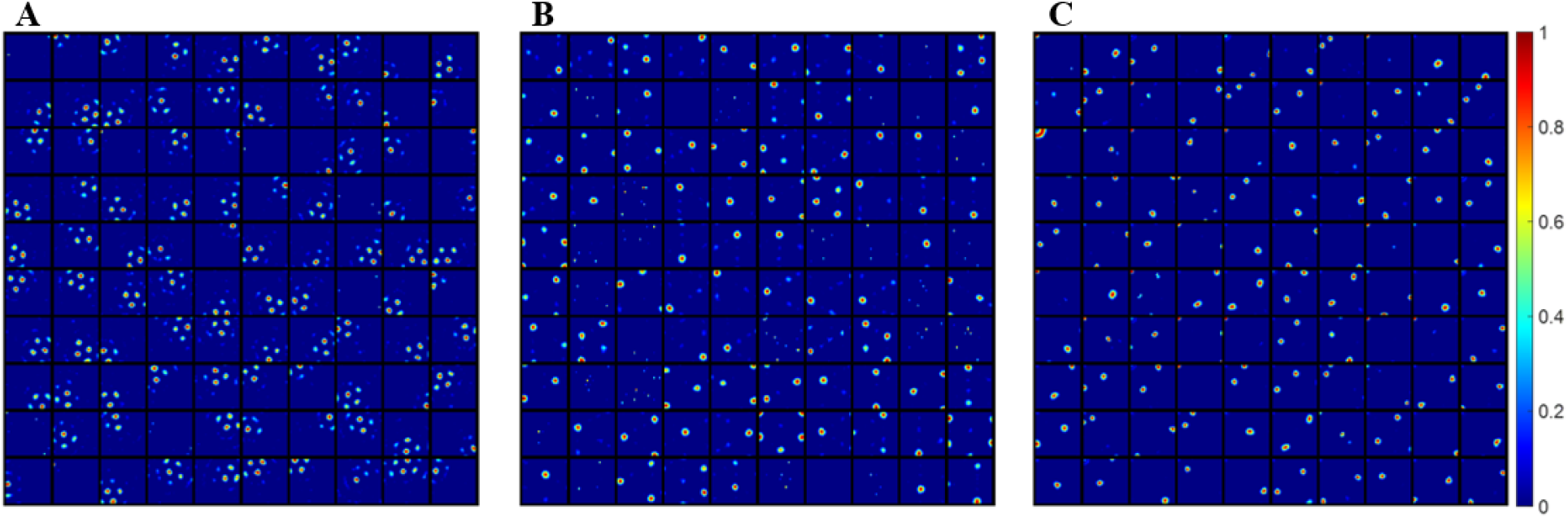
Modelled hippocampal cells with multiple firing locations start to emerge due to the lack of diversity in any of the grid parameters. Each block represents the firing field, **F** (Equation 10), of a modelled hippocampal cell in a 1m × 1m environment. Values in each block are normalised to [0 1]. **(A)** Lack of diverse grid spacings: 1 grid spacing, 6 orientations and 25 phases (*N*_*λ*_ = 1, *N*_*θ*_ = 6 and *N*_*x*_ = *N*_*y*_ = 5). **(B)** Lack of diverse grid orientations: 4 grid spacing, 1 orientation and 25 phases (*N*_*λ*_ = 4, *N*_*θ*_ = 1 and *N*_*x*_ = *N*_*y*_ = 5). **(C)** Lack of diverse grid phases: 4 grid spacing, 6 orientations and 1 phase (*N*_*λ*_ = 4, *N*_*θ*_ = 6 and *N*_*x*_ = *N*_*y*_ = 1).

Recall that the principle of sparse coding finds a linear representation of the input, namely the grid cell responses. Our results suggest that grid cells with less diversity in grid parameters are not sufficient to well represent the whole environment, so that the system gives an ambiguous representation of the spatial location. Therefore, the diverse grid cells found in the MEC are crucial to the emergence of hippocampal place cells if grid cells are the only entorhinal input to the hippocampus. The lack of diversity in afferent grid cells may be one possible factor explaining how cells with multiple firing locations emerge in the dentate gyrus.

#### When there are fewer modelled hippocampal cells

Simulations also show that a smaller number of modelled hippocampal cells, *N*_h_, can also cause the model to learn cells with multiple firing locations, even though the grid cells are diverse. This is illustrated in Figure 6, which shows the firing fields of modelled hippocampal cells when there are different numbers of modelled hippocampal cells in the network. The values of the remaining parameters are exactly the same as ones used in Figure 5 which shows well-learned place cells, except the number of modelled hippocampal cells, *N*_h_. Figure 6 demonstrates that as the number of modelled hippocampal cells, *N*_h_, decreases, cells with more firing-locations start to emerge. The less cells, the larger the proportion of cells with multiple firing locations that emerge. When *N*_h_ = 10 all modelled hippocampal cells have more than one firing location. When *N*_h_ = 20 there are five cells that are categorised as place cells. When *N*_h_ is larger than 30, almost all cells are found to have single-location firing fields.

**Figure 6:**
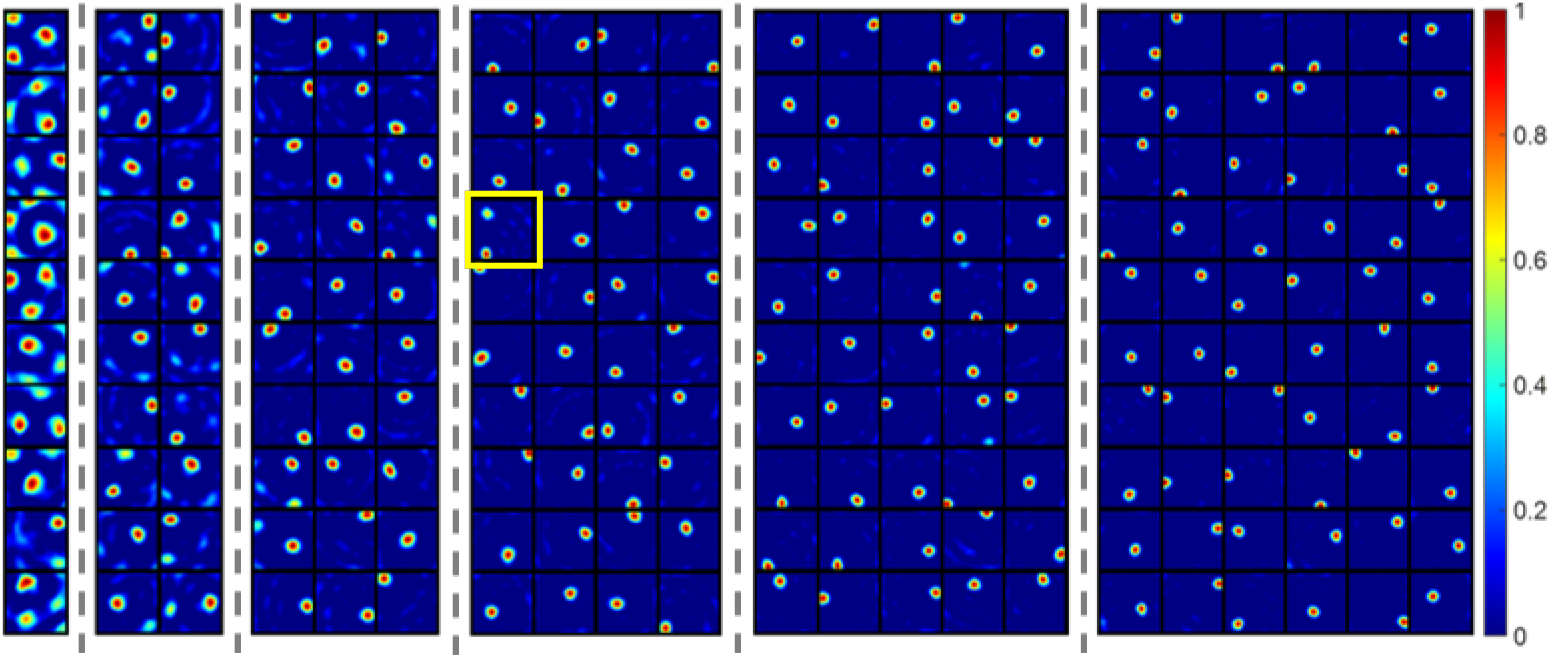
The firing fields, F (recovered by Equation 10), as the number of cells (*N*_h_) increases from 10 to 60 (separated by dash lines). A smaller number of cells, *N*_h_, leads to more cells with multiple firing fields after learning. The yellow box highlights the only cells with multiple firing locations when *N*_h_ *>*= 40. Each block represents the firing field of a cell in a 1m × 1m environment. Values in each block are normalised to [0 1]. Results shown in this figure have the same parameters as Figure 3 except the number of cells, *N*_h_.

Consequently, though a network with diverse grid cells can represent the spatial environment well, having less modelled hippocampal cells does not result in an unique representation of the spatial location with single-location place cells and the animal’s location might need to be encoded using more than 1 cell. This suggests more generally that hippocampal cells with multiple firing locations may be generated by having a small number of modelled hippocampal cells in the population that implements sparse coding.

### The spatial resolution of the hippocampal place map increases as more place cells are utilised to represent the environment

As discussed above, when *N*_h_ is larger than 30, almost all modelled hippocampal cells have a single-location firing field. In addition, the learned place cells tile the whole environment rather well with small values of the nearest distance (Equation 13). Furthermore, as *N*_h_ increases, the mean nearest distance and field breadth of the place field decreases (Figure 7) with relatively small variations, indicating that the spatial resolution of the neural representation by place cells improves.

**Figure 7:**
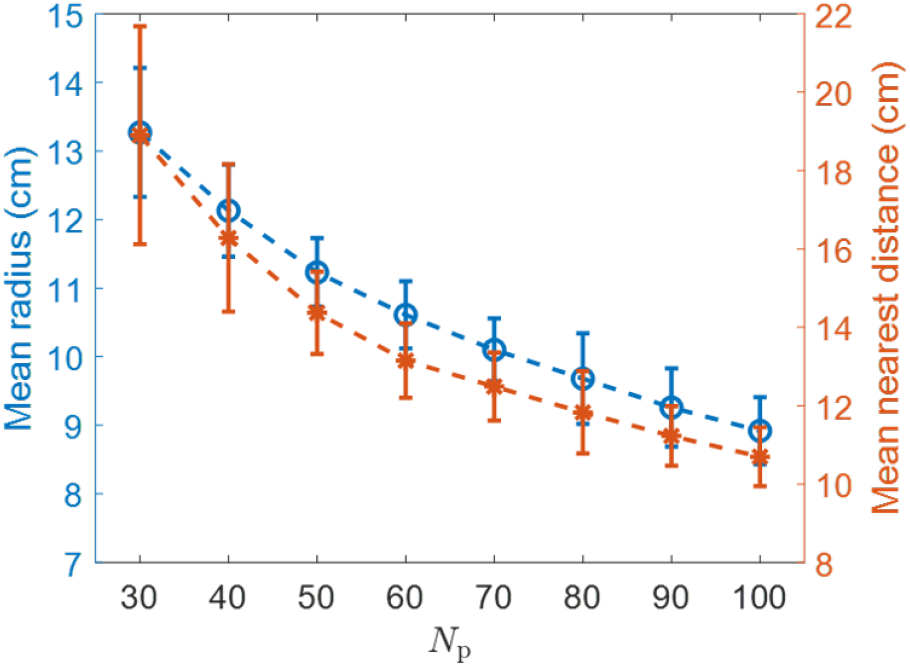
*N*_h_ vs. mean radius and *N*_h_ vs. mean nearest distance. As *N*_h_ increases, the mean radius and mean nearest distance decrease, indicating that a place map with higher spatial resolution is obtained. The error bars represent the standard deviations of radii and nearest distances.

### Model results are robust to realistic grid fields

When 600 more realistic grid cells sampled from four discrete grid modules (Materials and Methods) are used as the entorhinal input that incorporate the observed biological variability, the model can still learn a robust representation of the spatial location of the entire environment, as shown in Figure 8.

**Figure 8:**
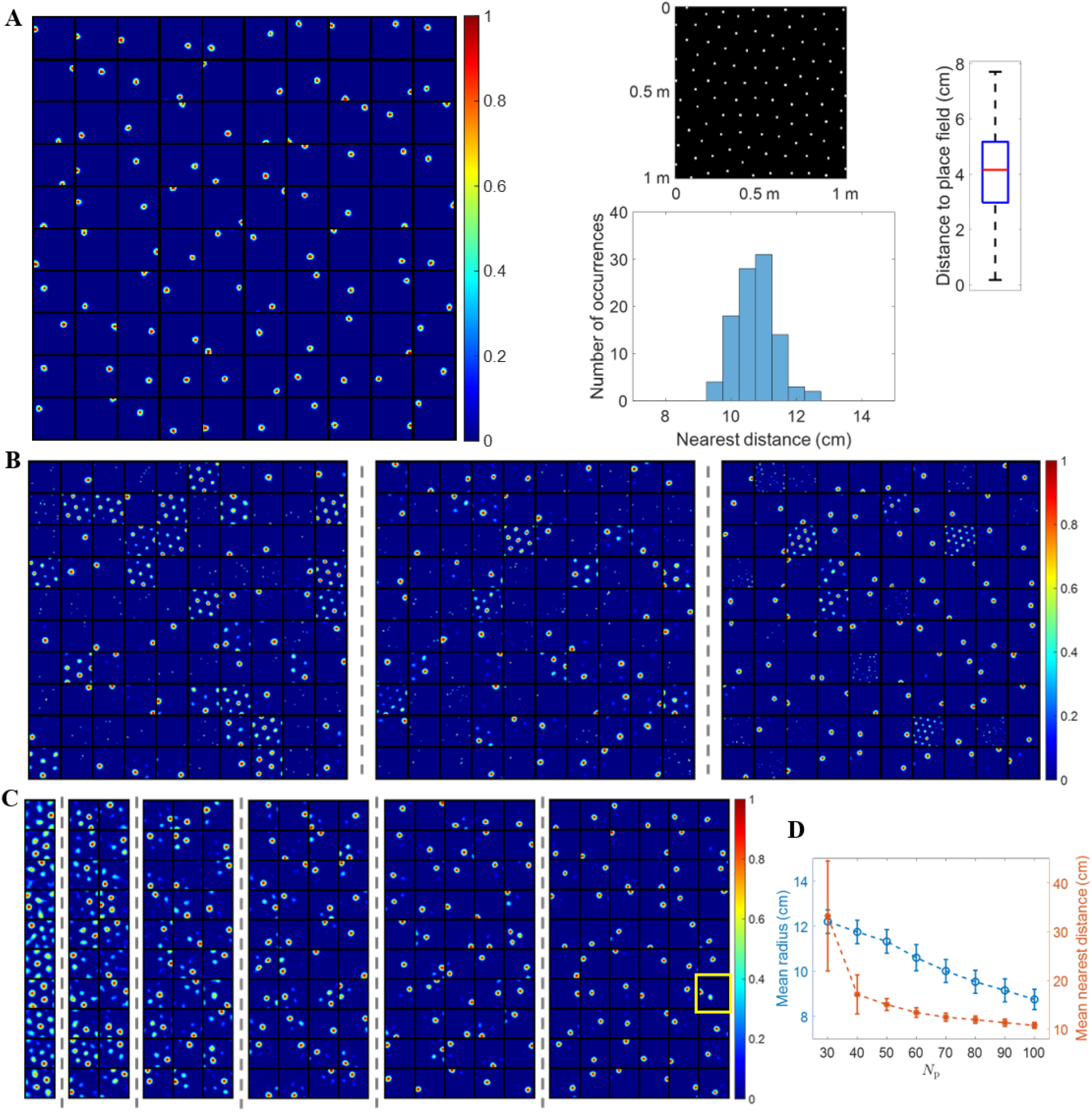
Results are robust when realistic grid fields are used. **(A)** Learned place cells evenly tile the entire spatial environment (similar to Figure 3). **(B)** The lack of diversity in grid spacing, orientation or phase (left, middle and right plot) leads to cells with multiple firing locations (similar to Figure 5). **(C)** The model with fewer modelled hippocampal cells leads to cells with multiple firing locations (similar to Figure 6). The yellow box highlights the only modelled hippocampal cell with multiple firing locations when *N*_h_ = 60. **(D)** Spatial resolution of the neural representation increases as *N*_h_ increases (similar to Figure 7). The error bars represent the standard deviations of radii and nearest distances.

Figure 8A shows that when grid fields are diverse in grid spacing, orientation and phase, each modelled hippocampal cell learns a single-location firing field such that centers of all place fields tile the entire spatial environment rather evenly. Furthermore, these place cells are positioned into a hexagonal pattern that packs the environment efficiently. The box plot of distance to place field shows fairly small values and indicates the whole environment is covered well. The distribution of nearest distance has mean 10.76 cm and standard deviation 0.62 cm, qualitatively consistent with results shown in Figure 3C (mean 10.70 cm and standard deviation 0.75 cm). Therefore, the learned place cells evenly tile the entire environment. Additionally, 600 grid cells sampled only from the two smallest grid modules are sufficient for place cells to emerge (data not shown).

Figure 8B shows that realistic grid fields with less diversity will cause the model to learn hippocampal cells with multiple firing locations. The left plot displays the firing fields of 100 modelled hippocampal cells with less diversity in grid spacing. The standard deviation of spacing in four grid modules is set to 0 cm instead of 8 cm while the standard deviation of orientation is still 3° and the phase is random. The middle plot shows firing fields when there is less diversity in grid orientation, where in each grid module the standard deviation of spacing is 4 cm, the standard deviation of orientation is 0° and phase is random. The right plot is for the case of less diversity in grid phase, where in each grid module the standard deviation of spacing is 8 cm, the standard deviation of orientation is 3° and phase is (0, 0) for all grid cells. Though the model learns hippocampal cells with multiple locations when there is less diversity in grid parameters, the learned place cells still tile the entire environment.

Figure 8C shows that having fewer modelled hippocampal cells causes the model to learn cells with multiple firing locations. The six plots separated by dashed lines in Figure 8C represent the firing fields of modelled hippocampal cells when the number of modelled hippocampal cells, *N*_h_, is 10, 20, 30, 40, 50, and 60, respectively. When *N*_h_ = 10 and 20, there is no place cell. When *N*_h_ ≥ 40, most modelled hippocampal cells are place cells. The yellow box indicates the only modelled hippocampal cell that is not a place cell when *N*_h_ = 60.

Similar to Figure 7, the neural representation of the spatial environment has better resolution (smaller radius and smaller nearest distance) as *N*_h_ increases, as seen from Figure 8D.

### The model can generate large place fields using grid cells

As discussed in a previous study (Neher et al., 2017), most existing models of place cells cannot produce large place fields, such as CA3 place cells with size around 1225 cm^2^. The model proposed here can generate large place fields by simply having grid cells with large grid spacings as the input to modelled hippocampal cells.

In this part of the study, more realistic grid cells are used as the entorhinal input to the hippocampus. In addition, only grid cells with grid spacings in the fourth module are used; i.e., the grid spacing is sampled from the normal distribution with mean 98.4 cm and standard deviation 8 cm, grid orientation is sampled from the distribution with mean 0° and standard deviation 3°, and grid phases are randomly chosen from a uniform distribution. Similarly, 600 grid cells are used. The number of modelled hippocampal cells, *N*_h_, is set to 20.

After learning, each modelled hippocampal cell pools a particular group of grid cells, similar to Figure 4A. 18 out of 20 modelled hippocampal cells satisfy the criteria of place cells defined in Materials and Methods. Figure 9A shows the 18 learned place cells. Figure 9C shows that these place cells have radii from 18.71 cm to 21.22 cm (mean 19.68 cm and standard deviation 0.75 cm). Therefore, the size of place fields ranges from 1099.76 cm^2^ to 1414.62 cm^2^. Figure 9B, D and E show that these 18 place cells with large size cover the entire environment rather evenly in a hexagonal pattern.

**Figure 9:**
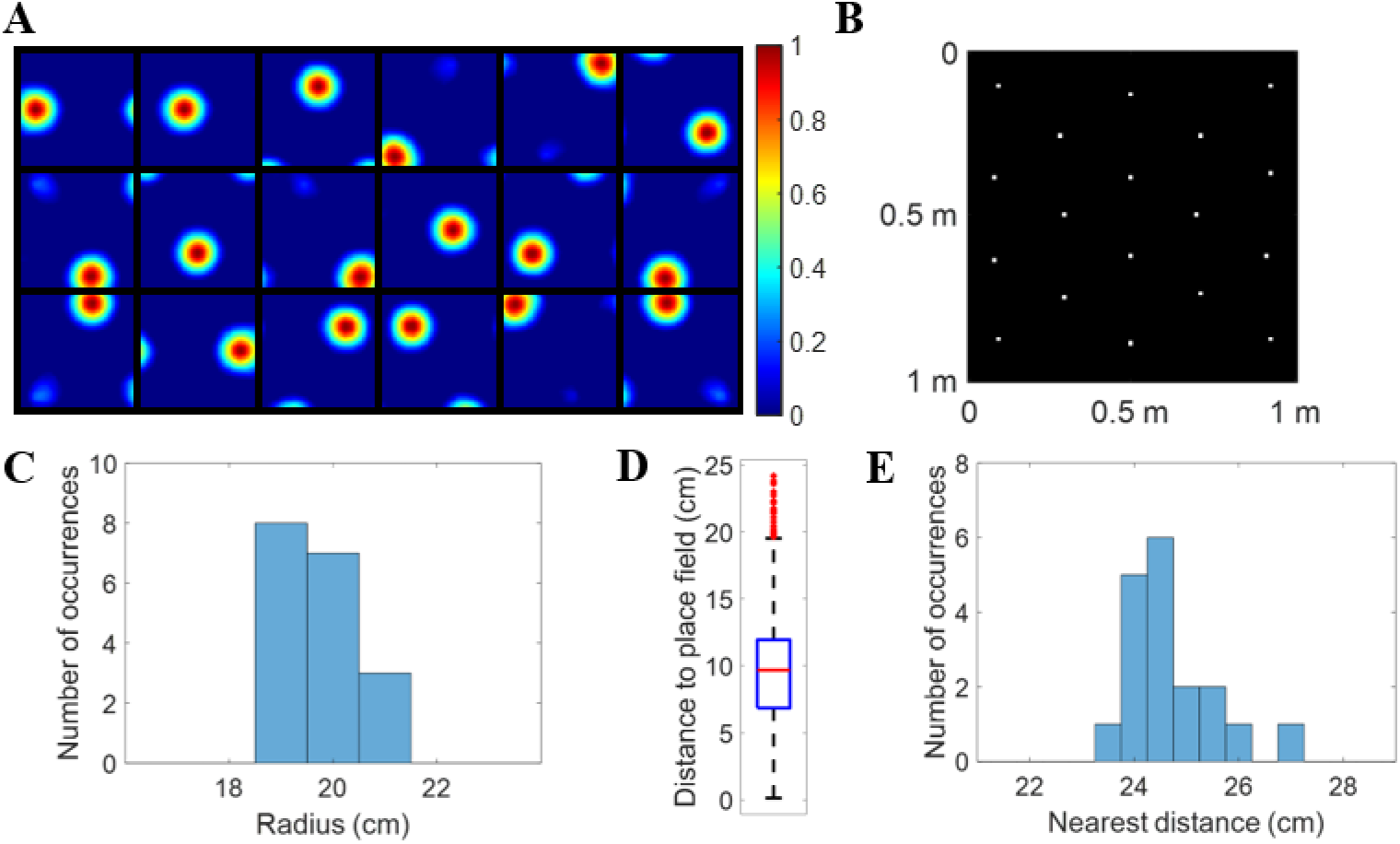
Large place fields emerge. **(A)** Firing fields, **F** (recovered by Equation 10), for 18 learned place cells. 18 out of 20 modelled hippocampal cells learn a place field at different spatial locations. Each block represents the firing field of a modelled hippocampal cell in a 1m × 1m environment. Values in each block are normalised to [0 1]. **(B)** Centers of place cells plotted together in the environment. Place cells evenly tile the whole spatial environment. **(C)** Histogram of radii for all place cells. **(D)** Box plot of distance to place field for all spatial locations in the spatial environment. The black lines at the bottom and top indicate the minimum and maximum, and the bottom edge of the blue box, red line inside the blue box and top edge of the blue box represent 25%, 50% (median) and 75% percentile of the data while outliers are excluded and represented as red dots. **(E)** Histogram of nearest distance for all place cells.

Above all, the model can learn large place cells if the afferent grid cells have large grid spacings, consistent with experimental evidence that the sizes of grid cells and place cells increase along the dorsal-ventral axis (Fyhn et al., 2007; Kjelstrup et al., 2008) and with topographic entorhinal-hippocampal projections along the dorsal-ventral axis (Dolorfo and Amaral, 1998).

### Weakly spatial cells in the EC are sufficient for hippocampal place cells to emerge

Recent experimental evidence shows that the emergence of hippocampal place cells happens earlier in development than grid cells (Langston et al., 2010; Wills et al., 2010). In addition, other experimental studies suggest that hippocampal place cells can still maintain their place fields even after grid cells are inactivated (Koenig et al., 2011; Brandon et al., 2014; Hales et al., 2014; Schlesiger et al., 2015). Though weakly spatial cells are more abundant than grid cells in the EC, how they contribute to the place fields of hippocampal place cells is still unclear. Here we show that even weakly spatial cells can provide sufficient spatial information for the emergence of place cells that have an accurate representation of spatial locations. This suggests that place cells can emerge throughout the development of MEC grid cells, from the initial weakly-tuned spatial pattern to the fully developed hexagonal grid pattern.

In this section, 600 weakly spatial cells are used as the entorhinal input to the hippocampus and there are 100 modelled hippocampal cells are used. We use a smaller learning rate (*η* = 0.01) and more epochs (30,000) for the learning here. The firing fields of ten example weakly spatial cells are shown in Figure 10A.

**Figure 10:**
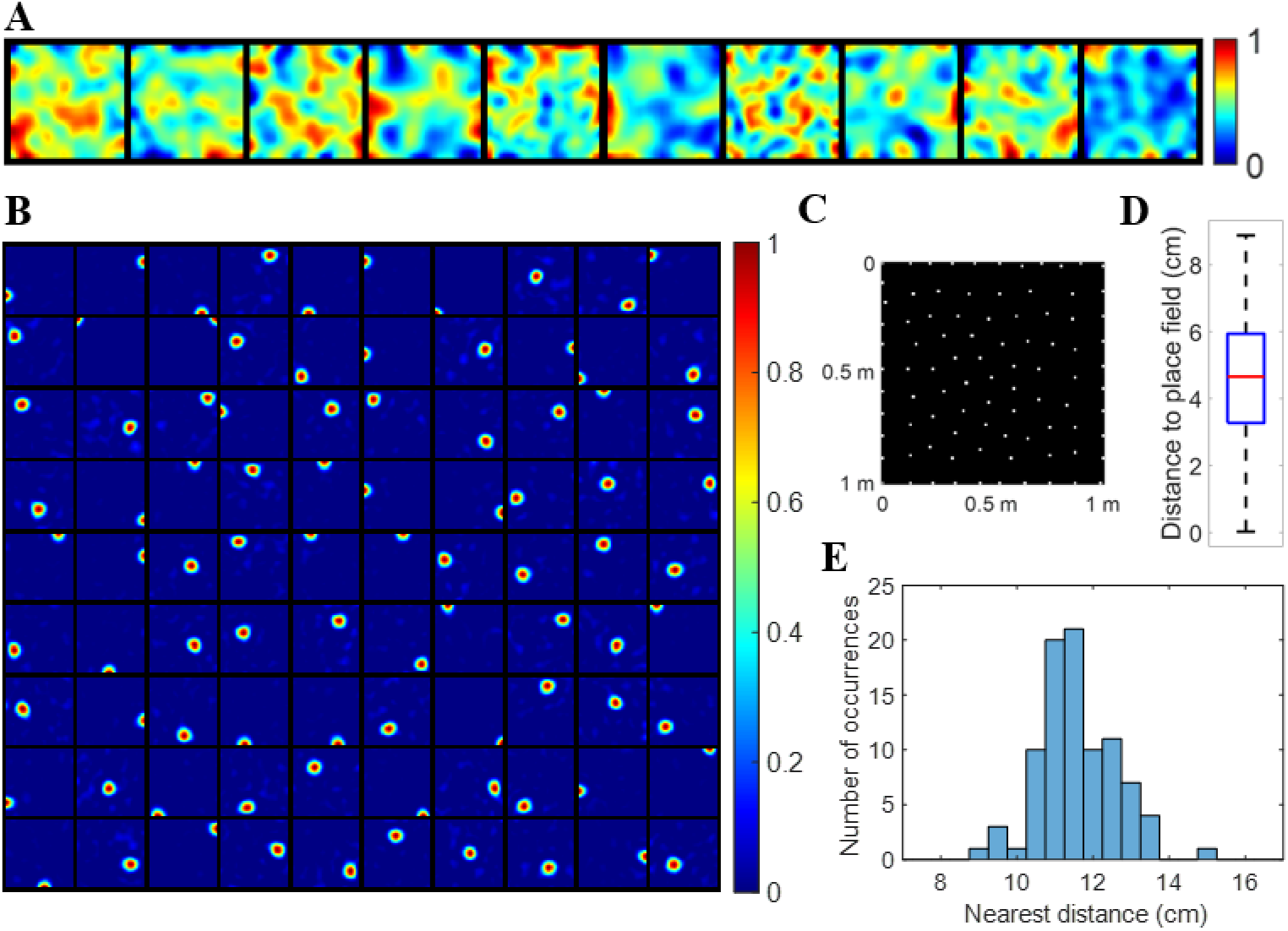
Weakly spatial cells are sufficient for place cells to emerge. **(A)** Examples of weakly spatial cells. Each block represents the firing field of a cell in a 1m × 1 m environment. **(B)** Firing fields, **F** (recovered by Equation 10), for 90 learned place cells. 90 out of 100 modelled hippocampal cells learn a place field at different spatial locations. Each block represents the firing field of a modelled hippocampal cell in a 1m × 1m environment. Values in each block are normalised to [0 1]. **(C)** Centers of 90 place cells plotted together in the environment. Place cells evenly tile the whole spatial environment with an efficient hexagonal pattern. **(D)** Box plot of distance to place field for all spatial locations in the spatial environment. The black lines at the bottom and top indicate the minimum and maximum, and the bottom edge of the blue box, red line inside the blue box and top edge of the blue box represent 25%, 50% (median) and 75% percentile of the data. In this box plot, there are no outliers. **(E)** Histogram of nearest distance for place cells.

#### An efficient hippocampal place map emerges

The firing fields of weakly spatial cells are very different from the periodic pattern of grid cells. Surprisingly, they can nevertheless provide sufficient spatial information such that the model based on sparse coding can decode entorhinal cell responses and give an accurate representation of the spatial location. Figure 10B shows the firing fields of learned place cells. Figure 10C, D and E show that the centers of place cells evenly tile the entire spatial environment with an efficient hexagonal pattern.

Compared with Figure 3 and 8A, using weakly spatial cells instead of grid cells results in learning a hippocampal place-map with less resolution. The mean radius of place fields using weakly spatial cells as the entorhinal input (Figure 10B: mean 11.45 cm with standard deviation 2.14 cm) is larger than Figure 3 and 8A (8.92 cm and 8.75 cm, respectively). Furthermore, the nearest distance in Figure 10E (mean 11.50 cm and standard deviation 0.94 cm) is also larger, compared with Figure 3 (mean 10.70 cm and standard deviation 0.75 cm) and Figure 8A (mean 10.76 cm and standard deviation 0.62 cm) when grid cells are used. The larger standard deviation in Figure 10E suggests that the irregular fields of weakly spatial cells lead to the less even tiling of place cells. However, the learned place map still covers the entire environment well with small distance to place field (Figure 10D) and efficiently with a hexagonal lattice (Figure 10C).

#### The learned place map is robust to noise even when weakly spatial cells are used

Furthermore, the model is quite robust to noise and an efficient place map can still be learned, even though a relatively strong noise is added to the modelled entorhinal cell responses in Equation 8:

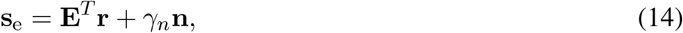

where **n** is the Gaussian noise with mean 0 and variance 1, and *γ*_*n*_ is the amplitude of the noise. Note that the maximal value of **E**^*T*^ **r** is 1 because **E** is normalised to have the maximum 1. We find that the model can still learn an efficient map when *γ*_*n*_ is 0.3 (Figure 11).

**Figure 11:**
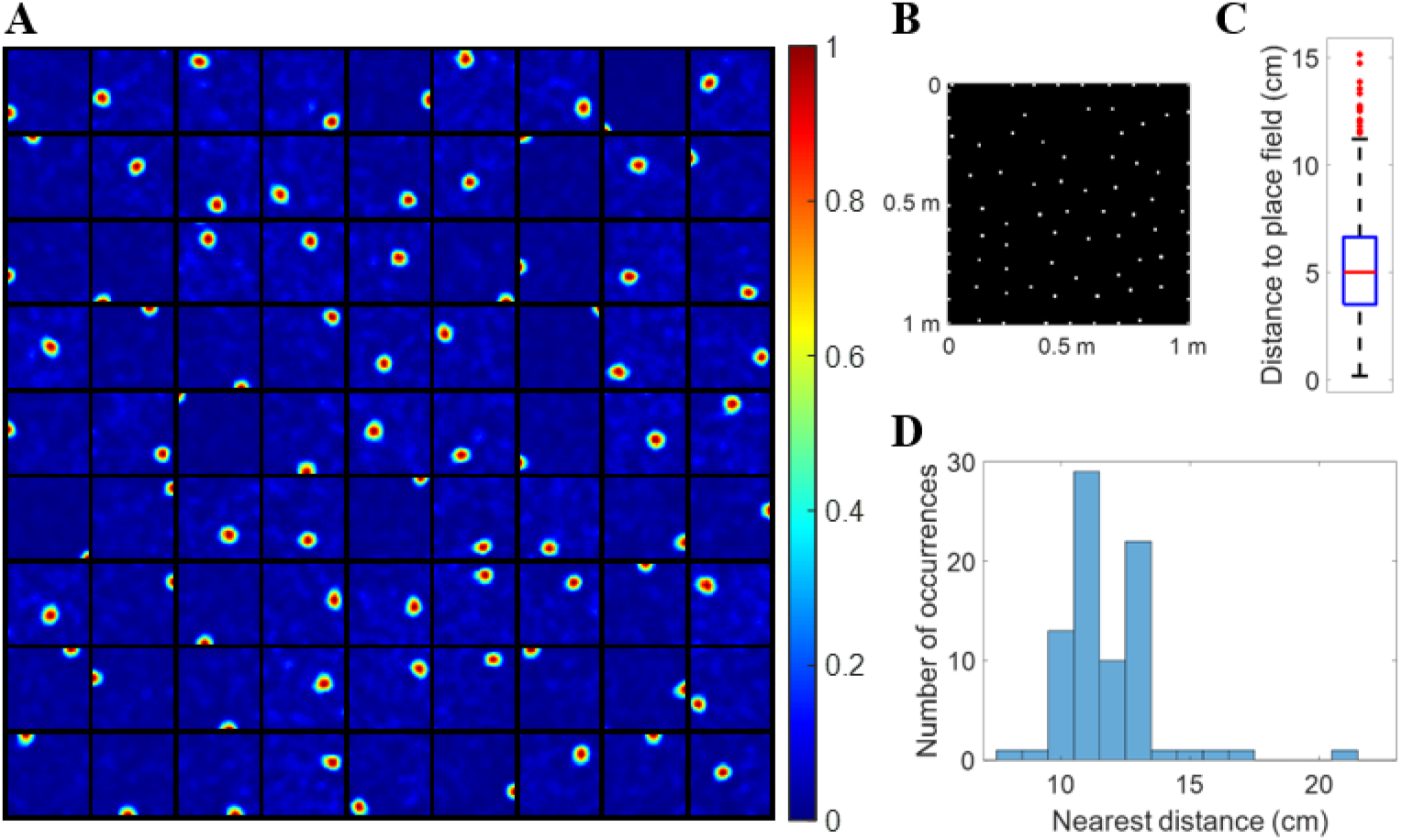
The model is robust to noise even when weakly spatial cells are used, *γ*_*n*_ = 0.3 in Equation 14. **(A)** Firing fields, **F** (recovered by Equation 10), for 80 learned place cells. 80 out of 100 modelled hippocampal cells learn a place field at different spatial locations. Each block represents the firing field of a modelled hippocampal cell in a 1m × 1m environment. Values in each block are normalised to [0 1]. **(B)** Centers of 80 place cells plotted together in the environment. Place cells tile the whole spatial environment. **(C)** Box plot of distance to place field for all spatial locations in the spatial environment. The black lines at the bottom and top indicate the minimum and maximum, and the bottom edge of the blue box, red line inside the blue box and top edge of the blue box represent 25%, 50% (median) and 75% percentile of the data while outliers are excluded and represented as red dots. **(D)** Histogram of nearest distance for place cells.

Though there are places in the environment not covered by place cells, generally the modelled hippocampal cells learn place fields that efficiently tile the entire environment.

Above all, the model is consistent with the experimental evidence that place cells emerge earlier than grid cells during development and a possible explanation is that the neural system can learn a hippocampal map even when the hexagonal spatial field is not well developed, and place cells can maintain their place fields when grid cells are inactivated because weakly spatial cells in the EC can lead to the emergence of a hippocampal place map.

Sparse coding can learn hippocampal place cells even though the input cells from the EC are weakly tuned to the spatial environment. Thus, input cells with stronger spatial selectivity can provide more spatial information so that unique place field can be decoded by sparse coding. Barry and Burgess (2007) used a learning model to learn place cells from responses of boundary vector cells that are selective to boundaries of the environment at particular angles and distances. Their result can be regarded as a special case of the results presented in this paper, where boundary vector cells are simply entorhinal input cells with stronger tuning of the spatial environment.

These weakly spatial cells can arise from any form of sensory inputs, such as visual input and auditory input, that encode spatial information. For example, the visual input at different locations of the environment actually carries information about spatial locations and consequently the afferent visual information to the EC can lead to weakly spatial cells. Moreover, the principle of sparse coding can cause EC cells to generate a hippocampal place map. The conjecture proposed here can explain a recent experimental study that shows that place cell firings mainly reflect visual inputs (Chen et al., 2019) and another experimental study that suggests homing abilities of mice even in darkness may not need accurate grid cell firing (Chen et al., 2016).

### The effect of environmental geometry on hippocampal cells may come from distorted entorhinal input

O’Keefe and Burgess (1996) found that place fields will be stretched following the environmental distortions. Later, Barry et al. (2007) found the same phenomenon for gird cells, namely that grid cells will be stretched along the distorted axis. Given the anatomical connection from the EC to hippocampus, it is natural to ask whether stretched place fields caused by the environment manipulation originate from stretched gird cells caused by the environment geometry. Our model shows that when grid fields are stretched along one axis of the environment, place fields will be stretched along that direction as well.

After the model is learned using 600 realistic grid cells as the entorhinal input to the hippocampus (Figure 8A), the entorhinal-hippocampal connection (**A**) is kept fixed. Then the spatial environment is changed to 2m × 1m; i.e., the environment is re-scaled along x-axis. In addition, gird fields are also stretched by a factor of 2 along x-axis. Next, we recover the firing fields of modelled hippocampal cells (described in Materials and Methods) with fixed connection and stretched grid cells. Figure 12 shows that modelled hippocampal cells are still selective to one location of the environment but their firing fields are stretched compared with the original place fields (left plot of Figure 8A).

**Figure 12:**
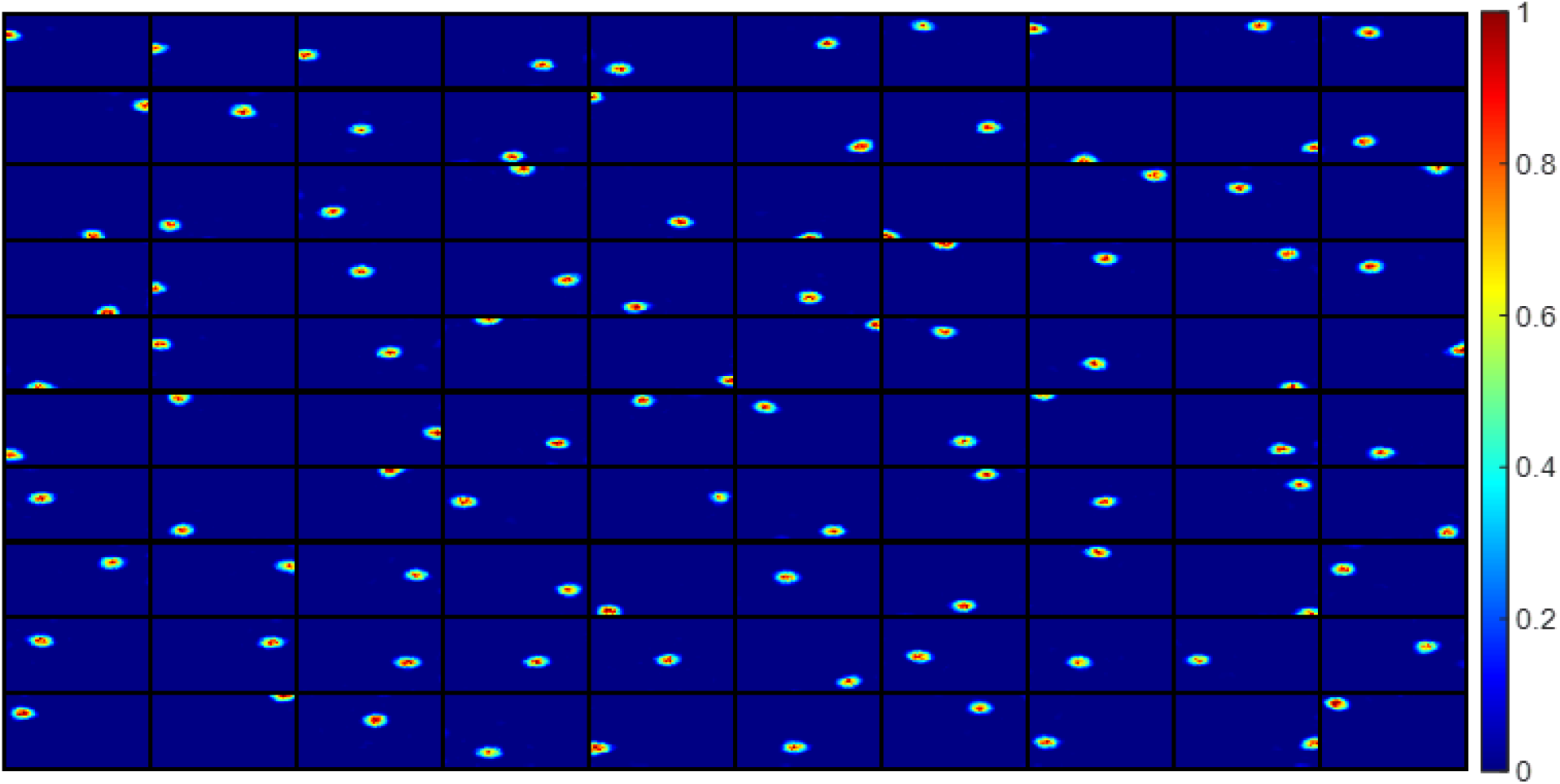
Firing fields of modelled hippocampal cells with distorted grid cells. Firing fields, **F** (recovered by Equation 10), for all 100 modelled hippocampal cells after the grid fields are stretched by a factor of 2 along the x-axis. Each block represents the firing field of a modelled hippocampal cell in a 2m × 1m environment. Values in each block are normalised to [0 1].

As grid cells are anchored to external landmarks (Hafting et al., 2005), grid fields are re-scaled by environmental changes (Barry et al., 2007). Our model suggests that the effect of environmental geometry on place cells may come from grid cells distorted by the environment manipulation. However, this does not rule out the possibility that the spatial selectivity of other entorhinal input to place cells is also altered by the environment, which causes the changes of place fields. If there is experimental data about the effect of environment geometry on other EC neurons that project to the hippocampus in the future, our model can be used to further explain what causes the change of hippocampal place fields.

### Navigation trajectory vs. random location

As described in Materials and Methods, the model is trained using random spatial locations uniformly sampled from the entire environment, so the spatial input to the model can uniformly cover the entire environment very quickly after some iterations. However, for a real navigation trajectory of a rat running freely in an environment, the route is continuous and there are spatial locations the rat has never been to. In this section, we used the same model as the one displayed in Figure 8A that has 600 realistic modelled grid cells as the entorhinal input to the model and 100 modelled hippocampal cells. Instead, the random spatial input to the model is replaced by a simulated running trajectory of a virtual rat. The trajectory is generated by a smoothed random walk using the method from D’Albis and Kempter (2017) with mean speed 0.25m/s and 20Hz of sampling positions. A simulated running trajectory of 3600 seconds is used to train the model. After learning, another simulated running trajectory of 1200 seconds is used to recover the firing fields of modelled hippocampal cells using the same method described in Equation 10.

Figure 13 shows the firing fields of 100 modelled hippocampal cells using the same realistic grid cells as Figure 8A but simulated running trajectory for training. Compared with Figure 8, firing fields trained and recovered by running trajectories are less circular due to non-uniform spatial locations used to train and recover the firing fields. Though the running trajectory of 3600s contains 72,000 spatial locations (much more than 20,000 spatial locations that are uniformly sampled to train the model in Figure 8), not all modelled hippocampal cells learn a place field. Positions along the running trajectory are continuous and are not sampled according to a uniform distribution, so the model has more data at some positions than others, causing the model to learn less place cells with less circular firing fields.

**Figure 13:**
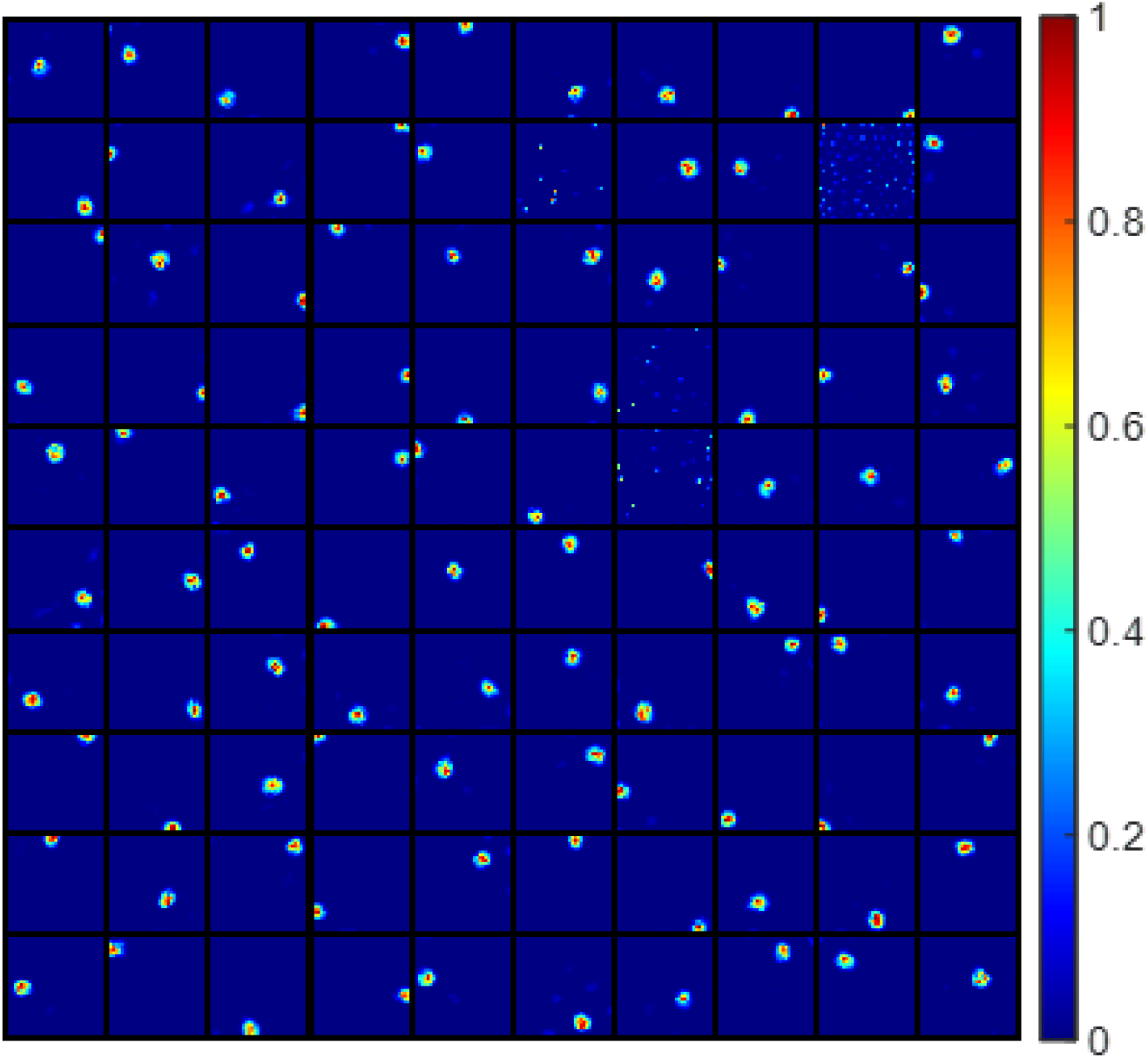
Firing fields of modelled hippocampal cells trained and recovered using simulated running trajectories. Firing fields, **F** (recovered by Equation 10), for all 100 modelled hippocampal cells. 4 out of 100 modelled hippocampal do not have an obvious firing field, while the other 96 modelled hippocampal cells learn place-like cells with one dominant firing location. Each block represents the firing field of a modelled hippocampal cell in a 1m×1m environment. Values in each block are normalised to [0 1].

## Discussion

### Summary

In this paper, we applied sparse coding with non-negative constraints to a hierarchical model of entorhinal-hippocampal network. Our results show that sparse coding can learn an efficient hippocampal place map that represents the entire environment when grid cells are diverse in grid spacing, orientation and phase. However, lack of diversity in grid cells or fewer modelled hippocampal cells leads to the emergence of cells with multiple firing locations, like those cells found in the dentate gyrus. In addition, weakly spatial cells in the EC are sufficient for sparse coding to learn hippocampal place cells.

### Comparison with other learning models

Our work differs significantly from previous studies on learning place cells from grid cell input (Rolls et al., 2006; Franzius et al., 2007b; Monaco and Abbott, 2011; Neher et al., 2017). Most importantly, our model investigates different types of entorhinal input: grid cells and weakly spatial cells. Also, among these learning models, we systematically investigate the influence of the diversity in grid cells using knowledge of grid modules (Stensola et al., 2012) upon the formation for hippocampal cells. Moreover, we demonstrate that learned hippocampal place cells can represent the entire spatial environment efficiently using a hexagonal lattice, consistent with a recent work in the conceptual state spaces (Mok and Love, 2019). Furthermore, the same model can produce cells with one firing location, multiple firing locations and large place field size, which can account for the emergence of a range of different observed hippocampal cell types. In addition, we demonstrate that weakly spatial cells in the EC can also provide sufficient spatial information for the emergence of the hippocampal place map after learning and the model is very robust to noise. Most importantly, all the results presented in this paper are generated by the same model, namely sparse coding with a non-negative constraint.

Though the principle of independent component analysis used by Franzius et al. (2007a,b) to learn place cells from grid cells is similar to the principle of sparse coding used here, the place cell examples in their paper are mostly near the boundary of the environment. However, our model can learn place cells at different places of the entire environment, generate a hippocampal place map that tiles the entire environment efficiently, and automatically provide the needed inhibition by the dynamics of sparse coding. One possible reason is that place cells in a population are not necessarily independent because nearby cells do overlap, so independent component analysis might put a too strong assumption. Also, the non-negativity introduced in this paper makes the model more similar to the real neural system, which might help the model uncover important biological properties.

### Properties of grid cells that are necessary for the emergence of place cells

Our model shows that the principle of sparse coding can learn an efficient place map using input from the EC, grid cells or weakly spatial cells. Though the model can learn place cells when only weakly spatial cells are used, it does not imply that grid cells are not necessary for the formation of place cells. Fiete et al. (2008) proposed that grid cells with different spacings and phases altogether form a residual system that efficiently encodes the spatial location. In addition, the triangular lattice of the grid pattern is known to be the solution to the optimal circle packing problem (Thue, 1892) and the geometric scale of grid spacings can represent the spatial environment efficiently (Wei et al., 2015). Our results are consistent with the concept that grid cells provide efficient information that can be retrieved to form a hippocampal place map. In our results, when 600 more realistic grid cells are used as the entorhinal input to hippocampal place cells, the ratio between the average of square of grid cell responses and the average of square of place cell responses is 0.31. However, this ratio becomes 0.84 when 600 weakly spatial cells are used as the entorhinal input, suggesting that grid cells are much more efficient for providing spatial information to the hippocampus. Another important property of grid cells that might be necessary for the emergence of place cells is the temporal property - phase precession. The experimental study conducted by Schlesiger et al. (2015) suggests that the MEC input to the hippocampus is necessary for the temporal property of hippocampal place cells. Therefore, though weakly spatial cells can provide sufficient spatial information to learn an efficient place map, grid cells that have phase precession might be required to account for the phase precession of hippocampal place cells. This is left for the future work.

### Non-grid cells in the entorhinal cortex

Though grid cells in the MEC have structured firing fields that lie on a hexagonal lattice, they only account for less than 1/3 of MEC cells and nearly 2/3 of MEC cells have spatial selectivity without well-characterised firing fields (Diehl et al., 2017). In the LEC, cells also display weak spatial specificity and convey nonspatial information (Hargreaves et al., 2005; Yoganarasimha et al., 2011). Our model explains how these upstream non-grid cells that contain weak spatial information can be used to form a hippocampal place map, which is consistent with increasingly more experimental evidence that suggests grid cells are not required to form the spatial firing field of place cells (Langston et al., 2010; Wills et al., 2010; Koenig et al., 2011; Brandon et al., 2014; Hales et al., 2014; Schlesiger et al., 2015).

Also, more recent studies have been investigating the effect of sensory input such as visual input on the firing of place cells. Chen and colleagues find that visual input largely determines place cells firing (Chen et al., 2013, 2016, 2019). In addition, a recent study shows that some neurons in V1 display location-specific firing (Haggerty and Ji, 2015). Therefore, we conjecture that neurons that respond to external stimuli in a spatial environment must contain spatial selectivity in any form and all these neurons contribute to the formation of a cognitive map. Entorhinal cortex, the gateway between the hippocampus and the neocortex of the brain, provides abundant information for the hippocampus and plays a crucial role here. Our results show that the principle of sparse coding can be one underlying principle that efficiently retrieves spatial information from the upstream. Furthermore, the efficient hexagonal pattern of the learned place map (Figures 3B, 8A, 9B and 10C) is consistent with the hexagonal clusters in a recent work in conceptual state spaces (Mok and Love, 2019), suggesting that the principle of sparse coding can be used to explain more functions of the hippocampus.

### Non-negativity

In recent computational studies of the navigational system of the brain, the concept of non-negativity is used several times. Dordek et al. (2016) applied non-negative principal component analysis to extract grid cells from place cell responses and the introduction of non-negativity pushes the model to converge to a hexagonal lattice from a square lattice when there is no non-negativity. D’Albis and Kempter (2017) build a single-cell spiking model for the grid cell and the non-linearity brought by the non-negative weights is sufficient for the emergence of a hexagonal lattice. Similarly, Sorscher et al. (2019) build a theory for the emergence of grid cells and the non-negativity constraint on the firing rates brings a symmetry-breaking effect that leads to the hexagonal firing fields. Unlike these studies that impose non-negativity into the model of grid cells, we use non-negative sparse coding to learn the hippocampal place cells. Non-negativity is introduced to account for some biological aspects such as non-negative firing rates and excitatory connection between cortical areas. In the original paper of non-negative sparse coding (Hoyer, 2003), the model learns Gabor-like V1 simple cells when trained on natural images; in other words, the non-negativity introduced is to account for the biological aspects and it is the principle of sparse coding that enables the model to learn V1 simple cells. The non-negative sparse coding used in this paper demonstrates that the principle of sparse coding can be used in the navigational system of the brain even when non-negativity constraints are incorporated. For our model, the non-negativity on neuronal responses is required to have a meaningful firing fields for the modelled hippocampal cells. However, when the non-negativity constraint on the connection **A** is removed, the modelled hippocampal cells still learn place fields that are similar to results demonstrated in this paper. Also, it is important to note that the non-negativity constraint on the entorhinal-hippocampal connection **A** does not prohibited the role of inhibition in the model. Instead, inhibition in the sparse coding model is vital to provide the competition needed to achieve the sparseness of the model. However, a very detailed model of how sparse coding can be implemented is still not very clear, which is discussed in the next section.

### Underlying neural circuits

Our study examines the extent to which sparse coding is as an underlying principle in the navigational system of the brain. However, the current model implies no specific neural circuits for the implementation of sparse coding, rather it is one of the principles that underlies the formation of the neural circuits. Neurophysiological and anatomical studies suggest that the EC and the hippocampus interact via a loop (Tamamaki and Nojyo, 1995; Tamamaki, 1997; Witter et al., 2014). Therefore, feedforward connections from the EC to the hippocampus, recurrent connections within the hippocampus, and feedback connections from the hippocampus to the EC all play an important role, though their specific contributions to the overall function of the network have not been fully uncovered yet. Renno’-Costa and Tort (2017) and Agmon and Burak (2020) investigated the coupling relationship between MEC grid cells and hippocampal place cells and showed that the proposed models can account for some experimental observations. However, how the underlying neural circuits can be implemented is still unclear. The proposed model based on sparse coding in this study does not rule out any of the network structures mentioned above, as sparse coding can be implemented in neural circuits either in a feedforward network with recurrent connections (Zylberberg et al., 2011) or a network with feedforward-feedback loops (Lian et al., 2019).

### Future work

The current study does not propose a specific biological neural circuit for implementing sparse coding in the entorhinal-hippocampal region, which is the study of ongoing work. Such a model of these neural circuits would need to take into account the experimentally known networks in this area. Also, other properties of place cells such as phase precession (O’Keefe and Recce, 1993), multiple place fields in large environments (Park et al., 2011) and place map in 3D environments (Grieves et al., 2020) will be investigated in the future. In addition, the model here used prefixed grid cells. We did not attempt here to provide a description for how grid cells emerge, but rather the grid cells are assumed to provide an efficient representation of the environment. It would be interesting to also investigate the role of sparse coding in how grid cells themselves emerge. It is hoped that such future work, which incorporates these aspects of the development process of both grid cells and place cells, will provide further insights into how the navigational system of the brain works. Sparse coding represents just one of a number of possible mechanisms that shape network structures, and much remains to be explored to incorporate other mechanisms, such as those associated with the complexities of metabotropic receptor effects, as discussed by Hasselmo et al. (2020).

## Acknowledgements

This work received funding from the Australian Government, via grant AUS-MURIB000001 associated with ONR MURI grant N00014-19-1-2571. We thank Kathrine Clarke, and Drs. Ali Almasi and Catherine Davey for helpful comments on the manuscript.

